# A preclinical pig model of Angelman syndrome mirrors the early developmental trajectory of the human condition

**DOI:** 10.1101/2025.03.04.641215

**Authors:** Luke S. Myers, Sarah G. Christian, Sean Simpson, Renan Sper, Clint Taylor, Laura Montes, Thomas B. C. Jepp, Daniela Ramos, Livia Schuller, Kranti Konganti, Wade Friedeck, Ozair Habib, McKaela Hodge, Alasdair J. Taylor, Ashley Coffell, Annalise Schlafer, Morgan Matt, Bradley Revell, Carol Knight, Cristina C. Barreña, FIRE consortium, William J. Murphy, Jorge Piedrahita, Scott V. Dindot

**Affiliations:** Department of Veterinary Pathobiology, College of Veterinary Medicine & Biomedical Sciences, Texas A&M University, College Station, TX, USA; School of Biosciences and Medicine, University of Surrey, Guildford, UK; Department of Molecular Biomedical Sciences, College of Veterinary Medicine, North Carolina State University, Raleigh, NC, USA; Comparative Medicine Institute, North Carolina State University, Raleigh, NC, USA; Interdisciplinary Graduate Program in Genetics and Genomics, Texas A&M University, College Station, TX, USA; Texas A&M University Institute for Genome Sciences and Society, Texas A&M University, College Station, TX, USA; Veterinary Medical Teaching Hospital, College of Veterinary Medicine & Biomedical Sciences, Texas A&M University, College Station, TX, USA; Department of Veterinary Integrative Biosciences, College of Veterinary Medicine & Biomedical Sciences, Texas A&M University, College Station, TX, USA; Ultragenyx Pharmaceutical Inc., Novato, CA, USA

**Keywords:** Angelman syndrome, UBE3A, imprinting, *Sus scrofa*, pig, CRISPR/Cas9

## Abstract

Angelman syndrome is a neurodevelopmental disorder characterized by severe motor and cognitive deficits. It is caused by the loss of the maternally inherited allele of the imprinted ubiquitin-protein ligase E3A (*UBE3A*) gene. Rodent models of Angelman syndrome do not fully recapitulate all the symptoms associated with the condition and are limited as a preclinical model for therapeutic development. Here, we show that pigs (*Sus scrofa*) with a maternally inherited deletion of *UBE3A* (*UBE3A*^−/+^) have altered postnatal behaviors, impaired vocalizations, reduced brain growth, motor incoordination, and ataxia. Neonatal *UBE3A*^−/+^ pigs exhibited several symptoms observed in infants with Angelman syndrome, including hypotonia, suckling deficits, and failure to thrive. Collectively, these findings are consistent with the pathophysiology and developmental trajectory observed in individuals with Angelman syndrome. We anticipate that this pig model will advance our understanding of the pathophysiology of Angelman syndrome and be used as a preclinical large animal model for therapeutic development.

## INTRODUCTION

Angelman syndrome is a devastating neurogenetic disorder with an incidence estimated at 1 in 12,000 to 24,000 live births^(1, 2)^. It is caused by the loss of expression or function of the maternal allele of the ubiquitin-protein ligase E3A (*UBE3A*) gene on human chromosome 15q11.2^(3)^. The loss of maternal *UBE3A* arises through different mechanisms, including large (5-7 Mb) *de novo* deletions of the 15q11.2-q13 region containing *UBE3A* (70-75%), *UBE3A* mutations (5-10%), imprinting center defects (2-5%), and paternal uniparental disomy (2-3%)^(4)^. Infants with Angelman syndrome behave typically at birth but frequently develop feeding difficulties due to poor suckling and problems swallowing^(3, 5–7)^. The first clinical sign of the disorder is developmental delay, which typically manifests at six months to one year of age^(7, 8)^. Motor incoordination, speech impairment, intellectual disability, and a unique happy demeanor are evident in infants and young children (1-3 years of age) and persist into adulthood^(7, 8)^. Seizures, abnormal electroencephalogram (EEG), sleep disorders, and microcephaly are also present in most but not all individuals^(7, 9, 10)^. There is currently no approved treatment specific to Angelman syndrome, but several disease-modifying and symptomatic-based treatments are under investigation^(11)^.

The *UBE3A* gene is subject to genomic imprinting in neurons of the central nervous system (CNS), a naturally occurring phenomenon in which the maternal allele is expressed, but the paternal allele is repressed^(12, 13)^. Imprinting of the paternal *UBE3A* allele is regulated by the *UBE3A* antisense (*UBE3A-AS*) transcript, which represents the distal end of the small nucleolar host gene 14 (*SNHG14*), an imprinted, paternally expressed, polycistronic transcription unit located downstream and in the opposite orientation to *UBE3A*. The distal end of *SNHG14*/*UBE3A-AS* is only expressed in CNS neurons, resulting in neuron-specific imprinting of *UBE3A*. In all other cell types and tissues, *UBE3A* is expressed from both parental alleles^(11, 14)^. The UBE3A protein functions as an E3 ubiquitin ligase and nuclear hormone transcriptional coactivator, with roles in the cell cycle, synaptic plasticity, synapse development, and cellular protein levels^(15)^. A noncoding isoform of *Ube3a* has also been shown to function as a microRNA sponge in the rat brain^(16)^. A complete loss of UBE3A expression in CNS neurons is thought to dysregulate critical pathways involved in synaptic function and plasticity, although the molecular pathogenesis underlying these deficits is poorly understood^(17)^.

Mouse and, more recently, rat models have played a vital role in Angelman syndrome research but have several limitations. The penetrance and expressivity of most phenotypes in the Angelman mouse models are strain and age-dependent and attenuated relative to those observed in patients^(18–33)^. Studies involving neonatal and juvenile rodents are also challenging because of their small size and altricial development. Importantly, translational research involving rodents frequently does not translate to patients, particularly for neurodevelopmental disorders^(34–37)^.

Pigs (*Sus scrofa*) have been used as animal models in biomedical research for centuries and are an ideal large animal model for studying CNS disorders in humans^(37–39)^. Compared to rodents, the pig brain is more similar to the human brain in developmental trajectory, size, structure, gyrencephalic folding, gene expression, neurotransmission, electrical activity, and white-to-gray matter proportion^(40–42)^. Pigs are precocial animals, enabling neonatal studies. They are also highly intelligent, capable of performing complex tasks. They vocalize using distinct sounds to convey emotional states and communicate with each other^(40, 43)^. Unlike other large animals, pigs have a short gestation period and produce multiple offspring, making them well-suited for generating sizable populations for experimentation^(37)^.

Here, we generated pigs with a complete deletion of the *UBE3A* gene using CRISPR/Cas9 and somatic cell nuclear transfer technologies. Pigs with a maternally inherited deletion of *UBE3A* recapitulated the core phenotypes associated with Angelman syndrome. Several phenotypes were penetrant at birth and more severe than those in rodent models.

## RESULTS

### Generation of a pig model of Angelman syndrome

We first examined the evolutionary conservation of the pig *UBE3A* gene with humans, cynomolgus macaques (*Macaca fascicularis*), mice, and rats. Analysis of multiple genome alignments spanning the *UBE3A* locus revealed long stretches of conserved sequences between humans, cynomolgus macaques, and pigs but shorter stretches between humans, mice, and rats (**Figure 1A**). Sequence alignments showed that the *UBE3A* genomic, mRNA, and amino acid sequences are more highly conserved between pigs and humans than between humans and mice and rats (**Figure 1B** and **Supplementary Table 1)**. Further analysis revealed that the rat and mouse *Ube3a* coding sequences are characterized by substantially higher (~2-fold) nucleotide substitution rates compared to humans and pigs (**Figure 1C**). Collectively, these findings show that the pig *UBE3A* gene shares greater sequence similarity with humans when compared to rodents, although rodents are phylogenetically closer to humans than pigs.

**Figure 1.**
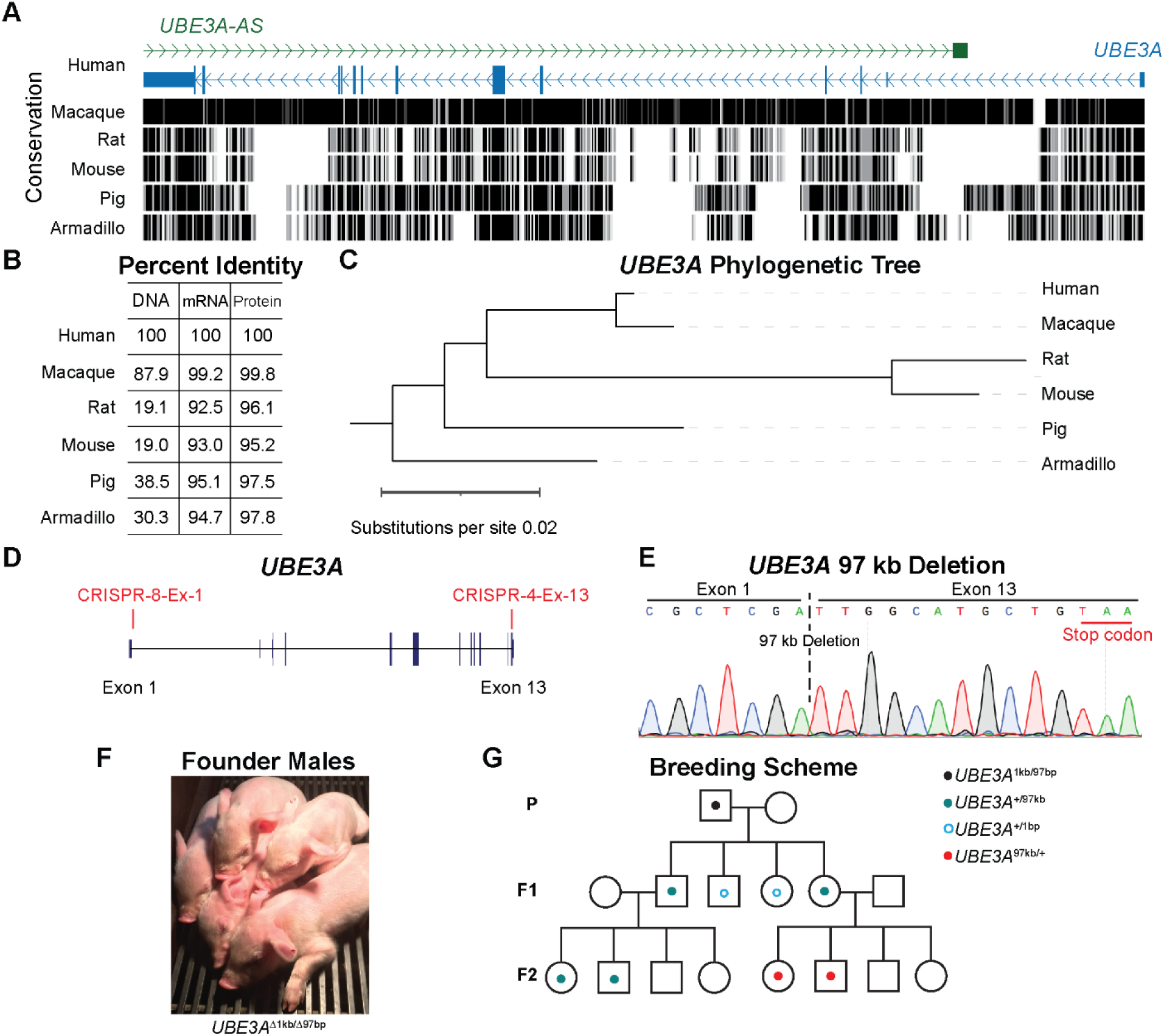
Generation of a pig model of Angelman syndrome. (**A**) Cactus alignment (Zoonomia project) between humans and the orthologous regions in cynomolgus macaques, rats, mice, pigs, and armadillos. (**B**) Percent identity of *UBE3A* DNA, mRNA, and protein sequences, relative to humans. (**C**) Phylogenetic tree illustrating the high substitution rate of *Ube3a* in rats and mice relative to other mammalian species. (**D**) Schematic illustrating the CRISPR-Cas9 target sites (red boxes) in exon 1 and exon 13 in the pig *UBE3A* gene. (**E**) Schematic of a Sanger sequencing chromatogram confirming the CRISPR/Cas9 mediated 97 kb deletion of *UBE3A*. (**F**) Image of founder male pigs generated by somatic cell nuclear transfer carrying a compound heterozygous deletion of *UBE3A* (*UBE3A*^97kb/1bp^). (**G**) Schematic of pedigree showing the breeding strategy to produce pigs with a maternal (*UBE3A*^−/+^) or paternal (*UBE3A*^+/−^) derived deletion of *UBE3A*.

To generate a pig model of Angelman syndrome, we first transfected CRISPR guide RNAs targeting the 5’ and 3’ ends of *UBE3A* (exons 1 and 13) into a fibroblast cell line derived from a male Yorkshire/Landrace pig. PCR screening of single-cell colonies identified five clonal cell lines harboring a complete (97 kb) deletion of *UBE3A* (**Figure 1D, Supplementary Table 2 and Supplementary Figure 1)**. Sanger sequencing confirmed the 97 kb deletion of the *UBE3A* gene on one allele and revealed a one bp deletion in exon 1 on the other allele (*UBE3A*^97kb/1bp^; **Figure 1E**). We then performed somatic cell nuclear transfer on cells from a single *UBE3A*^97kb/1bp^, resulting in two pregnancies producing seven viable males (**Figure 1F**). A single adult founder male (*UBE3A*^97kb/1bp^) was bred to wild-type (WT) females to generate F1 pigs with a paternally inherited deletion of *UBE3A*. Male and female F1 pigs carrying the 97 kb deletion of *UBE3A* were then bred to WT pigs to generate paternal *UBE3A*-deletion (*UBE3A^+/−^*) and maternal *UBE3A*-deletion (*UBE3A^−/+^*) pigs, respectively (**Figure 1G**). Whole-genome sequencing performed on a parent-offspring trio (WT male, *UBE3A^+/−^*female, and *UBE3A^−/+^* offspring) failed to identify any off-target mutations or evidence of donor plasmid integration into the genome (**Supplementary Data**). The *UBE3A*^97kb^ allele was transmitted in the expected Mendelian ratio in live births (*UBE3A^+/−^* n = 111 [49.1%], WT n = 115 [50.9%] [**Supplementary Table 3**]).

### The pig UBE3A gene is imprinted in the central nervous system

In humans and rodents, the *UBE3A*/*Ube3a* gene is only imprinted in the central nervous system (CNS)^(11–13, 44)^. To assess the imprinting status in pigs, we analyzed the expression of the parental *UBE3A* alleles in CNS and non-CNS tissues from WT, *UBE3A^+/−^*, and *UBE3A^−/+^* pigs (**Supplementary Table 4-6**). In CNS tissues, *UBE3A* RNA and protein expression was reduced by 70 - 85% in the *UBE3A^−/+^* pigs and 10 - 20% in the *UBE3A^+/−^*pigs relative to the WT pigs (**Figure 2A and 2B and Supplementary Figure 2A**), indicating the maternal *UBE3A* allele is preferentially expressed in the CNS. In non-CNS tissues, *UBE3A* RNA and protein expression was reduced by approximately 50% in both the *UBE3A^−/+^* and *UBE3A^+/−^* pigs relative to the WT pigs (**Figure 2A and 2B and Supplementary Figure 2A**), indicating the maternal and paternal *UBE3A* alleles are equally expressed in non-CNS tissues. The *UBE3A-AS* transcript was exclusively expressed in CNS tissues, consistent with the CNS-specific imprinting of *UBE3A* (**Supplementary Figure 2B)**. Lastly, immunohistochemical analysis showed that the UBE3A protein was enriched in the nuclei of cortical neurons in the WT pigs but was mostly undetectable in the *UBE3A^−/+^*pigs (**Figure 2C**). Collectively, these results show that the CNS-specific imprinting of *UBE3A* is conserved in pigs.

**Figure 2.**
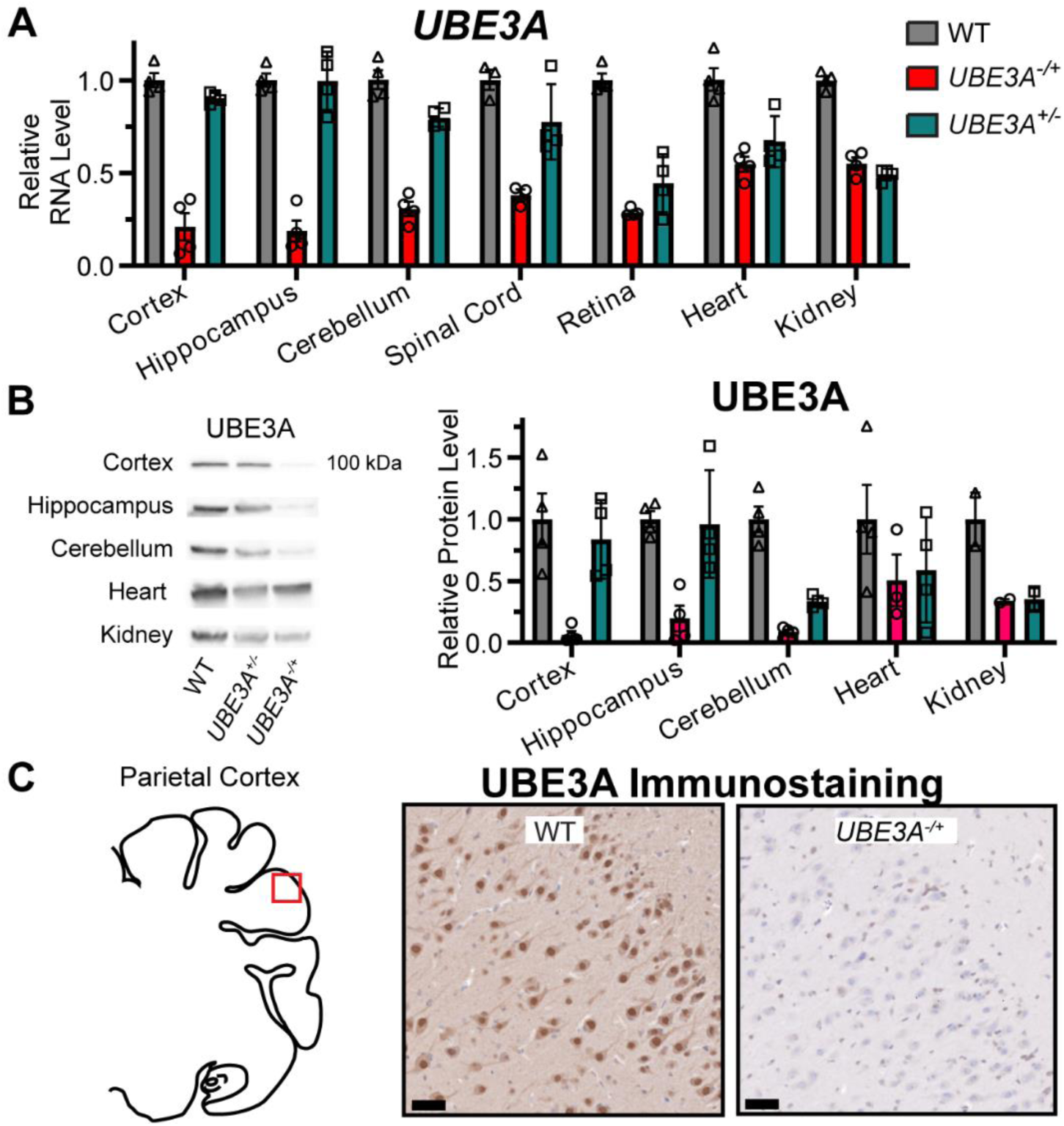
The pig *UBE3A* gene is imprinted in the central nervous system. (**A**) RT-PCR quantification of *UBE3A* RNA expression in WT, *UBE3A^−/+^*, and *UBE3A^+/−^*tissues, normalized to WT. Data presented as mean ± SEM. (**B**) Western blot quantification of UBE3A protein expression in WT, *UBE3A^−/+^*, and *UBE3A^+/−^* tissues, normalized to total protein and WT. Data presented as mean ± SEM. (**C**) Immunohistochemical analysis of UBE3A protein expression in the parietal cortex of WT and *UBE3A^−/+^*pigs; scale = 0.05 mm. Abbreviations: wild-type (WT), maternal *UBE3A* deletion (*UBE3A^−/+^*), and paternal *UBE3A* deletion (*UBE3A^+/−^*).

### UBE3A^−/+^ pigs have altered postnatal development

Newborns and infants with Angelman syndrome are often reported to have feeding difficulties (e.g., problems suckling and swallowing), hypotonia, and early signs of developmental delay (e.g., decreased babbling and crawling)^(3, 45)^. To determine whether postnatal development is affected in the *UBE3A^−/+^*pigs, we performed a series of studies to evaluate nursing, muscle tone, sensorimotor activity, and motor coordination across multiple cohorts of neonatal (0-28 days old), juvenile (28-70 days old), and adolescent (70-95 days old) pigs.

Cage-side observations of neonatal pigs revealed that in every litter at least one, often multiple, of the *UBE3A^−/+^* pigs had difficulty nursing, which appeared to stem from oral incoordination and the inability to establish a strong suckle (**Supplementary Video 1**). Significantly more *UBE3A^−/+^* pigs required supplemental feeding than their WT littermates (*UBE3A^−/+^*: 16/33 and WT: 2/38; p = 0.0004), often leading to failure to thrive. Using a modified limb resistance test on neonatal pigs after birth (0 – 3 days old) and after weaning (22 - 31 days old [**Supplementary Tables 4–6]**), we found that the newborn *UBE3A^−/+^* pigs had strikingly less muscle tone than the WT pigs (p = 0.0001 [**Figure 3A** and **Supplementary Figure 3A**]). Likewise, the post-weaned pigs had significantly less muscle tone than the WT pigs, although not as severe as the newborn pigs (**Figure 3A**).

**Figure 3.**
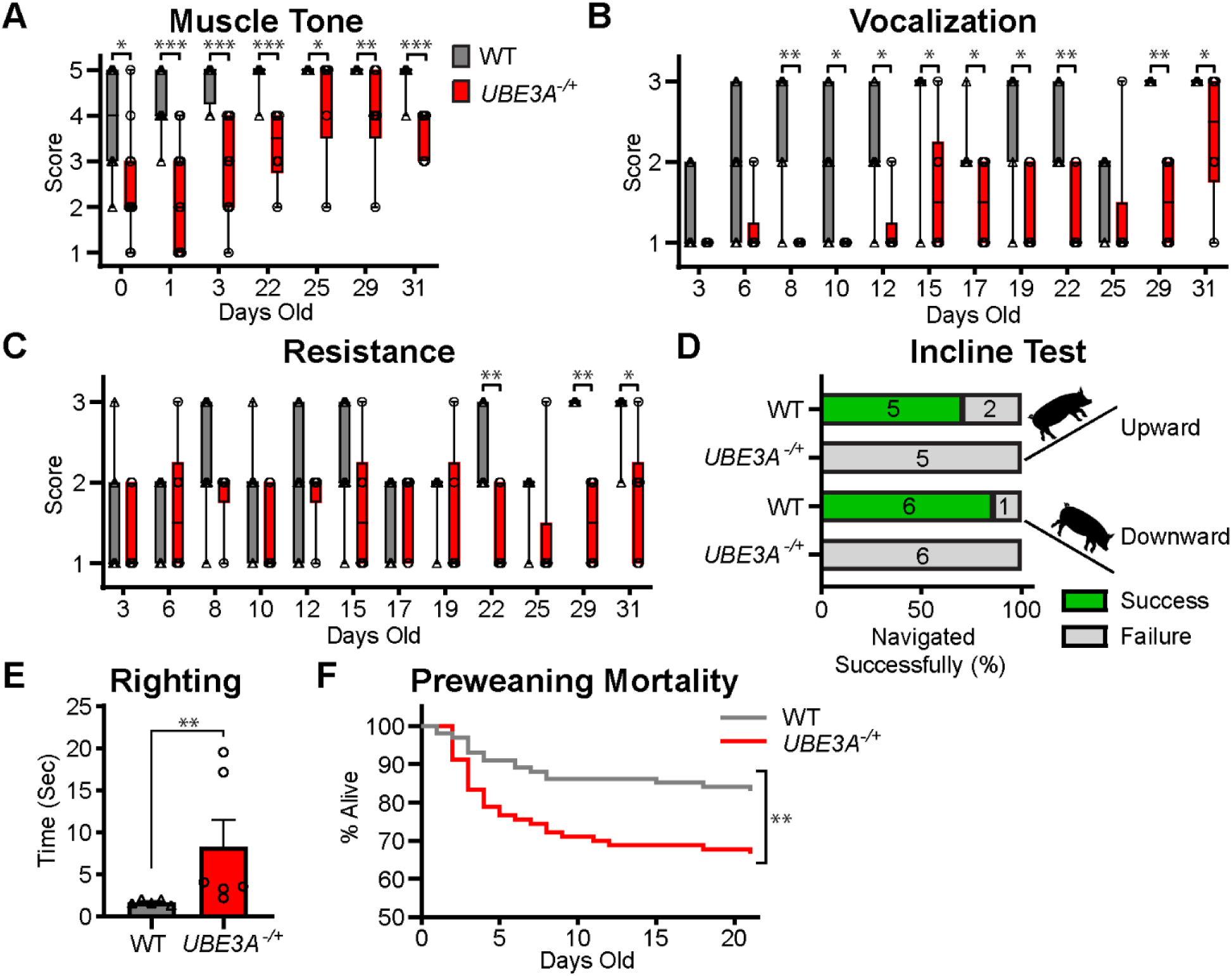
Impaired neonatal, juvenile, and adolescent development in *UBE3A^−/+^* pigs. (**A**) Muscle tone scores (1–5 scale) of WT and *UBE3A^−/+^*pigs. A score of 1 indicates low muscle tone with prominent shoulders, spine, and hips, and a score of 5 indicates high muscle tone with no visible rib or hip bones. Data distribution presented as a boxplot, Chi square ordinal logistic test. (**B**) Vocalization scores during the modified restraint test performed on WT and *UBE3A^−/+^* pigs (1: no vocalization; 2: grunting; 3: squealing). Data distribution presented as a boxplot, Ordinal Logistic Fit, FDR correction. (**C**) Resistance scores from the modified restraint test performed on WT and *UBE3A^−/+^* pigs (1: no movement; 2: leg kicking; 3: whole-body thrashing). Data distribution presented as a boxplot, Ordinal Logistic Fit, FDR correction. (**D**) Percent of neonatal (19-day old) WT and *UBE3A^−/+^*pigs successfully navigating the incline test. (**E**) Latency time for adolescent (71-day old) WT and *UBE3A^−/+^* pigs to transition from supine to standing in the righting reflex test. Data presented as mean ± SEM, Wilcoxon two sample exact test. (**F**) Kaplan-Meier survival analysis of WT and *UBE3A^−/+^* pigs, 2-tailed Fisher’s exact test. Abbreviations: wild-type (WT), maternal *UBE3A* deletion (*UBE3A^−/+^*), and paternal *UBE3A* deletion (*UBE3A^+/−^*). *\*P* < 0.05, \*\**P* < 0.01, and *\*\*\*P* < 0.0001.

We then performed a modified acute restraint stress test, which involves scoring a pig’s response (vocalizations and resistance) to mild restraint, to evaluate sensorimotor activity on multiple cohorts of neonatal and juvenile pigs (**Supplementary Video 2**). In the first cohort of neonatal pigs (3 to 31 days old; WT, n = 7; *UBE3A^−/+^*, n = 6 [**Supplementary Tables 4–6**]), the *UBE3A^−/+^* pigs vocalized significantly less than WT pigs at each age examined, except in newborn pigs (3-6 days old) and the first post-weaning assessment (25 days old) in which the WT pigs vocalized substantially less (**Figure 3B** and **Supplementary Table 7**). The resistance to restraint was also reduced in the *UBE3A^−/+^* pigs; however, significant differences were observed only in the post-weaned pigs (**Figure 3C** and **Supplementary Table 7**). A similar pattern was observed in a second cohort of neonatal and juvenile pigs (16 to 58 days old; WT, n = 5; *UBE3A^−/+^*, n = 5 [**Supplementary Tables 4–6**]) in which the *UBE3A^−/+^* pigs vocalized significantly less than WT pigs at each age, except for the first assessment after weaning (**Supplementary Figure 3B** and **Supplementary Table 7**). Most strikingly, many *UBE3A^−/+^* pigs never exhibited high-frequency vocalizations (i.e., squeals) during the testing, whereas the WT pigs were consistently highly vocal (**Figure 3B** and **Supplementary Figure 3B**). To determine whether these behavioral differences were due to the maternal *UBE3A* deletion, we assessed a cohort of neonatal and juvenile paternal *UBE3A* deletion pigs and their WT littermates (11 – 53 days old: *UBE3A^+/−^*, n = 5; WT, n = 9 [**Supplementary Tables 4–6**]). The behavioral responses of the *UBE3A^+/−^* pigs were indistinguishable from the WT pigs, demonstrating that the observed differences were specific to the maternal *UBE3A* deletion (**Supplementary Figure 3D-E**).

We next evaluated motor coordination using an incline test and a righting test. The incline test showed that neonatal *UBE3A^−/+^* pigs (19 days old: WT, n = 7 and *UBE3A^−/+^*, n = 6 [**Supplementary Tables 4–6**]) were unable to stay upright while navigating down the incline, in contrast to the WT pigs, regardless of their starting orientation (Upward: p = 0.005; Downward: p = 0.03 [**Figure 3D** and **Supplementary Figure 3F**]). Similarly, the righting test showed that adolescent *UBE3A^−/+^* pigs (71-74 days old: WT, n = 5 and *UBE3A^−/+^*, n = 6 [**Supplementary Tables 4–6**]) required significantly more time (approximately 5x longer) to stand than the WT pigs (p = 0.004 [**Figure 3E**]).

Lastly, we determined the survival rate of neonatal *UBE3A^−/+^*and WT pigs prior to weaning (21 days) across 21 litters (**Supplementary Tables 5–6**). Overall, the mortality rate of the *UBE3A^−/+^* pigs was twice as high as WT pigs (mortality rate: *UBE3A^−/+^*, 29/89 = 32.6%; WT, 15/101 =14.9%, p = 0.007 [**Figure 3F**]). The increased mortality rate of the *UBE3A^−/+^* pigs appeared to stem from a failure to thrive and sow-inflicted trauma during the first 12 days of life, after which time the survival rate of the *UBE3A^−/+^* pigs increased to 95%.

### UBE3A^−/+^ pigs vocalize less

Individuals with Angelman syndrome can emit audible utterances but have low expressive speech, and few can produce words or sentences^(46)^. To further assess vocalization in the *UBE3A^−/+^* pigs, we measured the total number of sounds (grunts and squeals) and the number of sounds by frequency (low-frequency grunt [0-8000 Hz], medium-frequency grunt [8001-12,500 Hz], and high-frequency squeal [12,501-22,000 Hz]) in neonatal, juvenile, and adolescent pigs (5 to 90 days old: WT, n = 16 and *UBE3A^−/+^*, n = 13 [**Supplementary Tables 4–6**]). Overall, the *UBE3A^−/+^* pigs produced significantly fewer sounds than the WT pigs at each age examined (**Figure 4A** and **Supplementary Table 7**). By frequency range, the *UBE3A^−/+^* pigs produced a similar number of low-frequency grunts (**Figure 4B**) but significantly fewer medium-frequency grunts (**Figure 4C**) and high-frequency squeals (**Figure 4D**). Visualization of the vocalization spectrograms (15 days old) revealed that the WT pigs emitted highly repetitive, uniformly structured low-frequency grunts, whereas the *UBE3A^−/+^* pigs emitted fewer, more irregular, non-uniform low-frequency grunts (**Figure 4E-F**). Together, these findings show that the *UBE3A^−/+^* pigs not only vocalize less but also have impaired vocalization structure.

**Figure 4.**
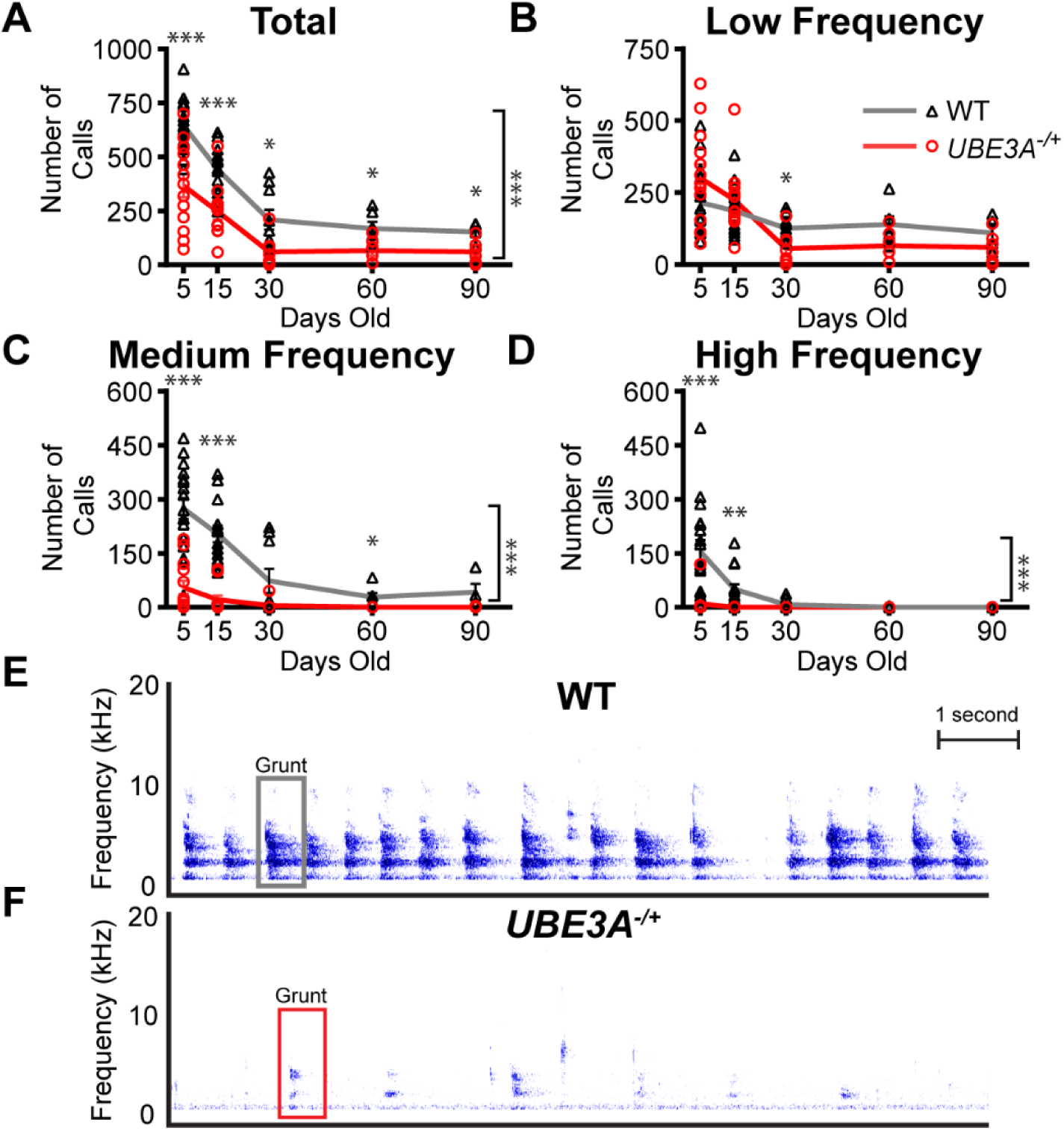
*UBE3A^−/+^* pigs vocalize less and have an abnormal vocalization structure. (**A**) Quantification of total vocalizations made by the WT and *UBE3A^−/+^* pigs during development. Data presented as mean ± SEM, Student’s t, all pairwise comparison. (**B**) Quantification of low-frequency grunts (0–8000 Hz) made by the WT and *UBE3A^−/+^*pigs. Data presented as mean ± SEM, Student’s t, all pairwise comparison. (**C**) Quantification of medium-frequency grunts (8001–12,500 Hz) made by the WT and *UBE3A^−/+^*pigs. Data presented as mean ± SEM, Student’s t, all pairwise comparison. (**D**) Quantification of high-frequency squeals (12,501–22,000 Hz) made by the WT and *UBE3A^−/+^* pigs. Data presented as mean ± SEM, Student’s t, all pairwise comparison. (**E and F**) Representative spectrograms of a 15-day old WT and *UBE3A^−/+^* pigs illustrate fewer vocalizations and abnormal vocalization structure in the *UBE3A^−/+^* pigs. Spectrograms display intensity and frequency over 10 seconds. Abbreviations: wild-type (WT) and maternal *UBE3A* deletion (*UBE3A^−/+^*). *\*P* < 0.05, \*\**P* < 0.01, and *\*\*\*P* < 0.0001.

### UBE3A^−/+^ pigs have an ataxic gait

Individuals with Angelman syndrome have an ataxic gait characterized by a wide stance and a short, highly variable stride length^(7, 47)^. To examine gait in the *UBE3A^−/+^* pigs, we measured the step width and stride length in juvenile pigs (33-44 days old: WT, n = 11 and *UBE3A^−/+^*, n = 11 [**Supplementary Figure 4**]). The *UBE3A^−/+^* pigs had a significantly wider step (forelimb: p = 0.02; and hindlimb: p = 0.01) and significantly shorter stride length (forelimb: p = 0.03; and hindlimb: p = 0.03) than the WT pigs (**Figure 5A–D**), indicating the *UBE3A^−/+^* pigs have an ataxic gait.

**Figure 5.**
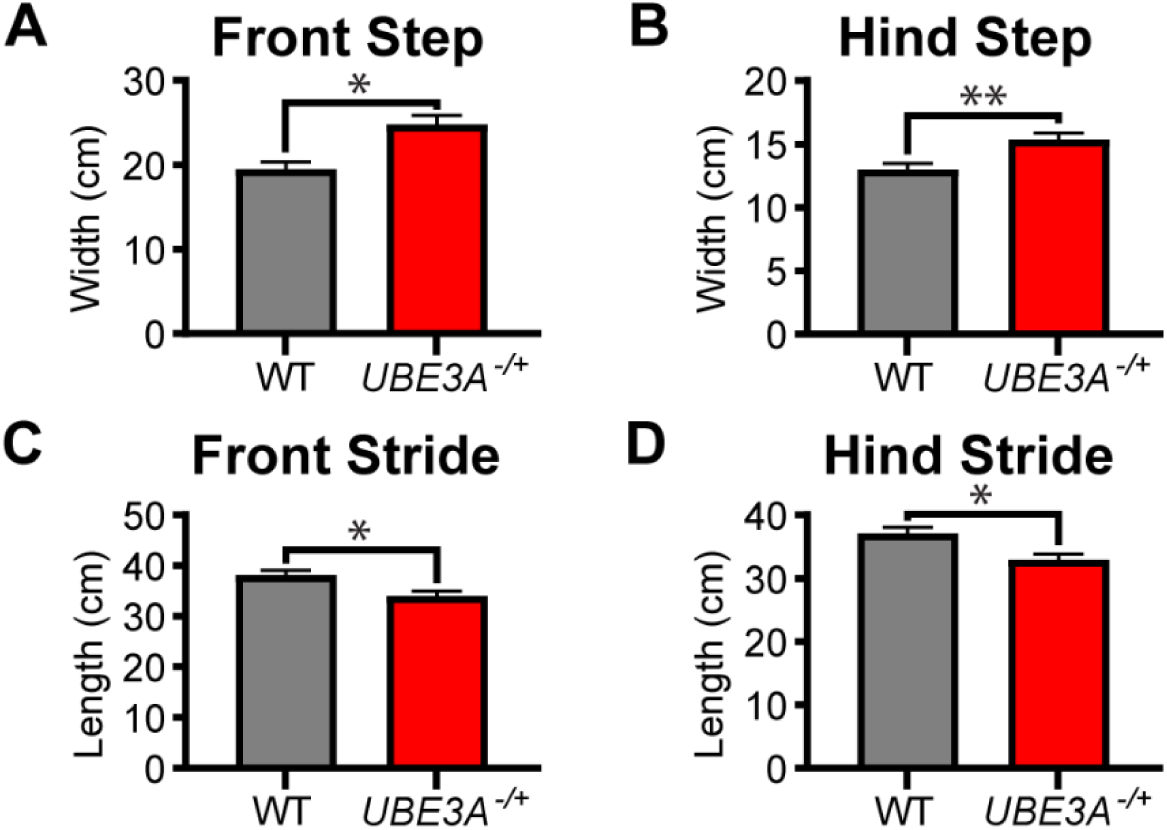
Juvenile *UBE3A^−/+^* pigs have an ataxic gait. (**A and B**) Gait map measurements of the front and hind step lengths and (**C and D**) front and hind stride lengths of juvenile (33-44 days old) WT and *UBE3A^−/+^* pigs. Data presented as mean ± SEM, Students t, all pairwise comparison. Abbreviations: wild-type (WT) and maternal *UBE3A* deletion (*UBE3A^−/+^*). *\*P* < 0.05 and *\*\*P* < 0.01.

### UBE3A^−/+^ pigs are hypoactive

Children with Angelman syndrome often exhibit hyperactive and hypermotoric behaviors, which tend to diminish with age^(48–51)^. Conversely, rodent models of Angelman syndrome are hypoactive, although these findings are limited to adult animals^(52, 53)^. To assess activity in the *UBE3A^−/+^* pigs, we used an open field arena to measure the distance traveled, time traveling, and velocity of pigs at different neonatal and adolescent stages (5, 15, 30, 60, and 90 days old: WT, n = 16 and *UBE3A^−/+^*, n = 13 [**Supplementary Tables 4–6** and **Supplementary Figure 5A**]). The distance traveled, percent time moving, and velocity of the neonatal *UBE3A^−/+^* pigs (5 and 15 days old) were significantly lower than the WT pigs (**Figure 6A-B**, **Supplementary Table 7, and Supplementary Figure 5B-C**), which was largely due to the lack of general exploratory behavior. The activity level of the older *UBE3A^−/+^*pigs was similar and not significantly different than the WT pigs (Distance traveled, F = 0.4; and cumulative movement, F = 0.9 [**Figure 6A-B**]), which was due to decreasing exploratory behavior in the WT animals (Distance traveled, F = 0.0001; and cumulative movement, F = 0.0001 [**Figure 6A-B**]).

**Figure 6.**
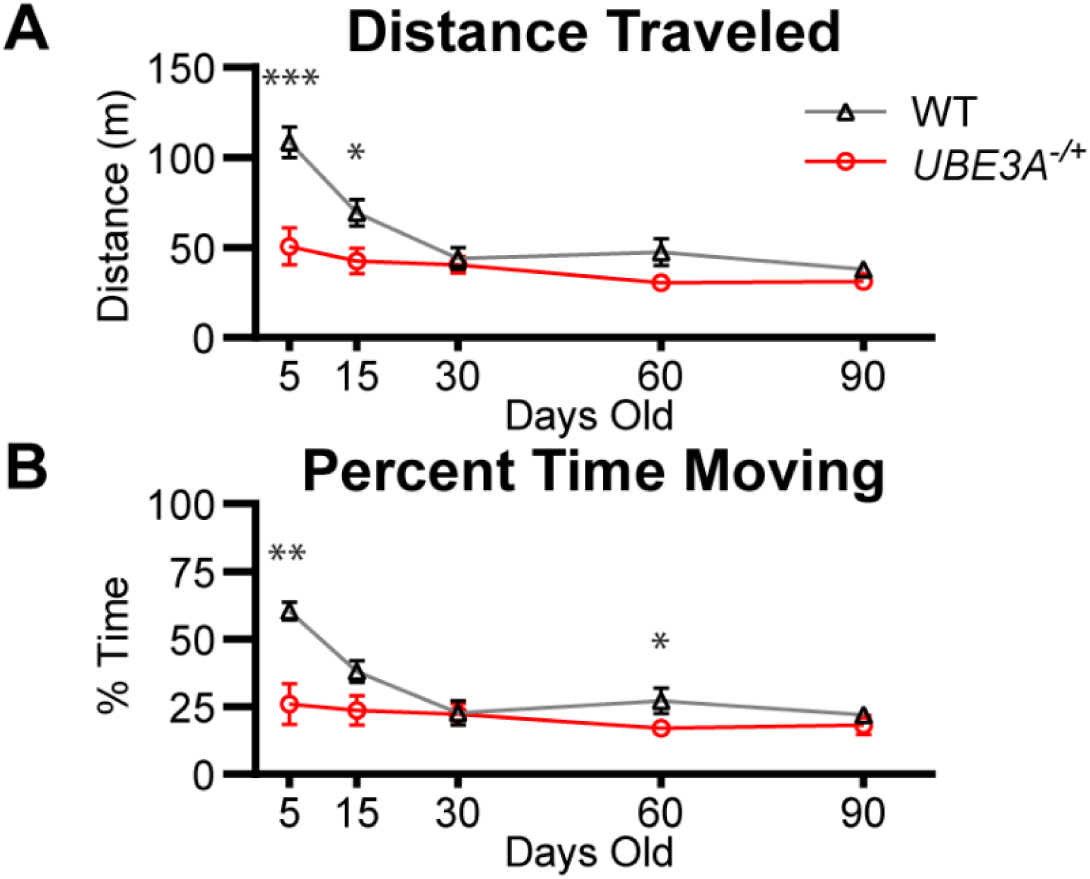
Neonatal and juvenile *UBE3A^−/+^* pigs are hypoactive. (**A and B**) Activity (distance traveled and percent time moving) of WT and *UBE3A^−/+^* pigs in an open field arena. Data presented as mean ± SEM, Wilcoxon two sample exact test. Abbreviations: wild-type (WT) and maternal *UBE3A* deletion (*UBE3A^−/+^*). *\*P* < 0.05, \*\**P* < 0.01, and *\*\*\*P* < 0.0001.

### UBE3A^−/+^ pigs have reduced body weight and brain size

Individuals with a *UBE3A* deletion exhibit lower birth weight^(10, 54)^; however, body weight and mean body mass index normalize in adults ^(55)^. In contrast, adult mice — but not rats — lacking a maternal *Ube3a* allele become overweight ^(21, 56)^. Progressive microcephaly is reported in 24–80% of individuals with Angelman syndrome^(9, 10)^, and brain development delays are similarly observed in both mouse and rat models^(31, 57, 58)^.

We first assessed body growth in neonatal, adolescent, and juvenile pigs (1–95 days old: WT, n = 94; *UBE3A^−/+^*, n = 66; *UBE3A^+/−^*, n = 13 [**Supplementary Tables 5–6**]). The *UBE3A*^−/+^ pigs weighed significantly less than WT pigs at every age except the final time point (**Figure 7A** and **Supplementary Table 7**). At 1–3 days old, *UBE3A^−/+^* pigs weighed ~22% less than WT pigs, and by 55–65 days old, they weighed ~12% less. However, despite their lower weight, they followed a similar growth trajectory as WT pigs. *UBE3A^+/−^*pigs displayed an intermediate weight pattern, generally falling between the WT and *UBE3A^−/+^* pigs, except at 55–65 days old, when their weight aligned more closely with *UBE3A^−/+^* pigs (**Figure 7A**).

**Figure 7.**
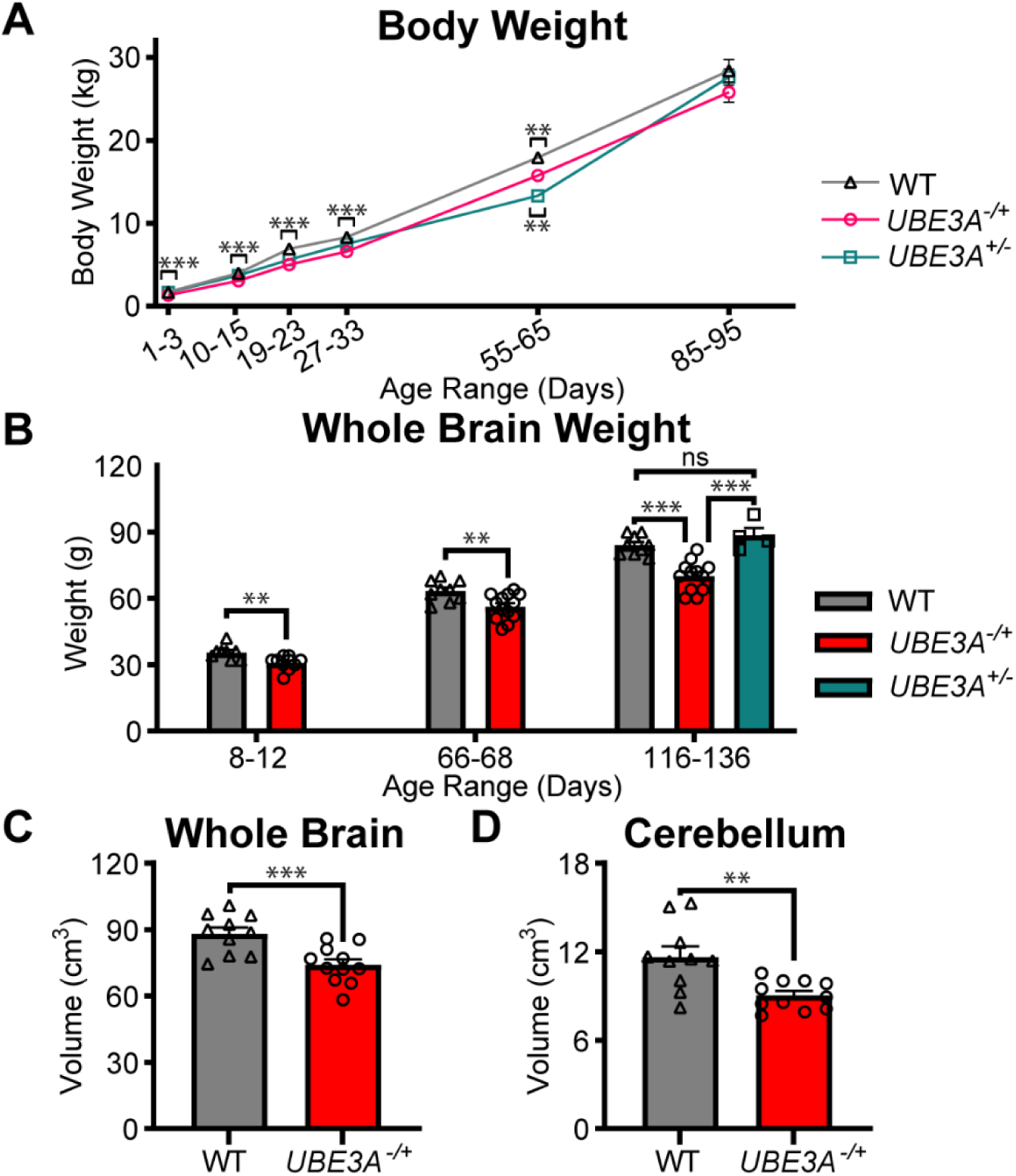
*UBE3A^−/+^* pigs have reduced body weight and brain volume. (**A**) Longitudinal analysis of body weight during neonatal, juvenile, and adolescent development. Data presented as mean ± SEM, all pairwise comparisons - Tukey HSD. (**B**) Gross brain weights of the WT and *UBE3A^−/+^*at different age ranges during postnatal development, including the brain weights of the *UBE3A^+/−^*pigs at 116–136 days old. Data presented as mean ± SEM, ANOVA Tukey-Kramer HSD. (**C and D**) Postmortem MRI analysis of whole brain and cerebellum volumes of adolescent (96-115 days old) WT and *UBE3A^−/+^* pigs. Data presented as mean ± SEM, Students t, all pairwise comparison. Abbreviations: wild-type (WT), maternal *UBE3A* deletion (*UBE3A^−/+^*), and paternal *UBE3A* deletion (*UBE3A^+/−^*); *\*\*P* < 0.01 and *\*\*\*P* < 0.0001; ns, not significant.

We then examined brain development in neonatal, juvenile, and adolescent *UBE3A^−/+^* pigs (8-12 days old; WT, n = 8 and *UBE3A^−/+^*, n = 10; 65-66 days old; WT, n = 9 and *UBE3A^−/+^*, n = 13; and, 116-136 days old; WT, n = 9, *UBE3A^−/+^*, n = 12, and *UBE3A^+/−^*, n = 4 [**Supplementary Tables 4–6**]). The mean gross brain weights of the *UBE3A^−/+^* pigs were significantly less than the WT pigs at each developmental stage examined, with the mean brain weights reduced by 13% at 8-12, 11% at 65-66, and 17% at 116-136 days old (**Figure 7B** and **Supplementary Table 7**). To determine if the reduced brain growth is due to the loss of the maternal *UBE3A* allele, we also determined the gross brain weights of adolescent *UBE3A^+/−^* pigs (116-130 days old). The brain weights of the *UBE3A^+/−^* pigs were similar and not significantly different from the WT pigs (p = 0.45) but significantly more than *UBE3A^−/+^* pigs (p = 0.0001; **Figure 7B**). Collectively, these findings indicate that brain development is reduced in the *UBE3A^−/+^*pigs and that the reduced growth is due to the loss of the maternal — but not paternal — *UBE3A* allele. Lastly, we used magnetic resonance imaging to measure post-mortem brain and cerebellar volume in adolescent pigs (97-115 days old: WT, n = 10 and *UBE3A^−/+^*, n = 11 [**Supplementary Tables 4–6**]). The brain and cerebellar volumes in the *UBE3A^−/+^* pigs were significantly less than the WT pigs, with mean volumes reduced by 16.1% and 22.3%, respectively (brain, p = 0.001; and cerebellum, p = 0.003 [**Figure 7C-D**]).

## DISCUSSION

We generated a pig model of Angelman syndrome using CRISPR/Cas9 and somatic cell nuclear transfer technologies. Our findings show that pigs with a deletion of the maternal — but not paternal — *UBE3A* allele mirror many early developmental phenotypes displayed by individuals with Angelman syndrome, including nursing problems, hypotonia, motor deficits, impaired vocalization, and delayed brain growth^(3, 5–10)^. These behavioral phenotypes were highly penetrant among animals across multiple litters and were more severe than comparable phenotypes observed in rodent models of Angelman syndrome, such as motor deficits and impaired vocalization.

Although mice and rats are phylogenetically closer to humans than pigs, we show that the pig *UBE3A* gene (genomic, coding, and amino acid sequences) is more conserved with humans than mice and rats, further supporting our prior finding that the *Ube3a* locus in murid rodents has diverged substantially from other placental mammals^(11)^. Our findings that the pig *UBE3A* gene is imprinted in the CNS but not in peripheral organs align with the neuron-specific imprinted expression of the *UBE3A* gene, further indicating that imprinting of *UBE3A* is a highly conserved regulatory mechanism among placental mammals^(11–14)^.

The neonatal *UBE3A*^−/+^ pigs have nursing problems, hypotonia, motor deficits, and abnormal sensorimotor responses, consistent with the early developmental symptoms observed in neonates and infants with Angelman syndrome^(7)^. These findings underscore the utility of using pigs as an animal model, as studies involving neonatal rodents are limited by their altricial development. Hypotonia, reduced vocalizations, and motor deficits were evident in almost all neonatal *UBE3A*^−/+^ pigs. These deficits were persistent throughout development and strikingly different from those of their wild-type littermates. The nursing problems observed in the neonatal *UBE3A*^−/+^ pigs, however, were highly variable within and across litters. This finding is likely due to the inherent variability in this phenotype, which is thought to stem from hypotonia of the cleft palate^(7)^, and our inability to evaluate nursing behavior objectively. Nevertheless, many neonatal *UBE3A^−/+^*pigs failed to thrive, as evidenced by the necessary interventional feeding and increased mortality rate. Interestingly, the mortality rate normalized after the pigs reached ~2 weeks of age, suggesting the *UBE3A^−/+^* pigs can survive beyond this critical period.

Impaired vocalization was one of the most striking and penetrant phenotypes observed in the *UBE3A^−/+^* pigs. The *UBE3A^−/+^*pigs vocalized remarkably less than WT pigs at every age, including adults, and in every behavioral context. For example, in addition to the quantitative assessment of vocalizations presented here, we observed that the *UBE3A^−/+^*pigs failed to emit squeals before being fed, which is a robust associative learning behavior. Whether this atypical behavior is due to deficits in cognition (e.g., learning and memory) or vocalization (e.g., motor control or hypotonia), or both, requires further investigation. Additionally, our results show that not only did the *UBE3A^−/+^*pigs vocalize less but they also struggled with the complexity of vocal sound production, as evidenced by the abnormal spectrograms. Overall, these findings closely mirror the communication deficits in human patients^(7, 46)^, providing a robust behavioral outcome measure for testing preclinical therapies.

Our findings that the *UBE3A^−/+^* pigs have motor deficits are consistent with those observed in individuals with Angelman syndrome^(7, 47)^. In addition to our gait analysis, we observed that the *UBE3A^−/+^* pigs often displayed other motor deficits, such as problems pivoting, walking off a step (e.g., out of the pen), and lying down. They were also noted to walk with a stilted gait, a phenotype frequently observed in individuals with Angelman syndrome^(7, 47)^. Whether these phenotypes are due to generalized hypotonia, motor neuron deficits, or a combination of both warrants further investigation. Nevertheless, the motor problems observed in the *UBE3A^−/+^* pigs were noticeable.

We acknowledge several limitations of the study. First, although our experiments included animals from multiple litters and produced consistent findings, some analyses lacked sufficient animals to assess sex as a biological factor with confidence (e.g., incline test). Second, our phenotypic analysis was not comprehensive. Additional studies are needed to evaluate other disease-relevant phenotypes, such as learning and memory, brain electrical activity (EEG), seizure susceptibility, synapse development, and molecular alterations.

In conclusion, we generated and characterized a pig model of Angelman syndrome. This is the first large animal model of the syndrome and one of the first large animal models of a monogenic neurodevelopmental disorder. Overall, our study highlights the profound developmental impact of maternal *UBE3A* deficiency from birth and aligns with clinical observations in human infants^(7)^, reinforcing the validity of using this pig model to study the early developmental trajectory of Angelman syndrome. The use of pigs also provides several advantages, such as the ability to perform longitudinal studies on a gyrencephalic brain with developmental milestones similar to those of humans^(40–42)^. This model also enables exploration of the molecular mechanisms underlying the disorder and opens promising avenues for developing and testing therapeutic interventions, allowing for assessment of the broader neurological and behavioral deficits observed in humans with this condition. The anatomical and physiological similarities between pigs and humans make this model highly suitable for pharmacokinetic and pharmacodynamic studies, further enhancing its utility in preclinical drug development. Future studies should focus on therapeutic development, biomarker discovery, and longitudinal assessments of cognitive and motor functions. Although additional characterization of this model is needed, our findings suggest that this pig model will be a valuable resource for translational research in Angelman syndrome.

## EXPERIMENTAL MODEL AND SUBJECT DETAILS

Experiments on all live animals were conducted under the guidance, supervision, and approval of the North Carolina State University or Texas A&M University Institutional Animal Care and Use Committees.

### Pig Housing and Experimental Design

Pigs were housed in climate-controlled rooms with age-appropriate temperature and light cycles. Feed was provided ad libitum until day 40, then adjusted to 2–4% of body weight daily. Animals were group-housed, except when single housing was required for treatments. Behavioral studies were conducted between 10:00 a.m. and 4:00 p.m. to avoid confounding factors from feeding schedules, with pigs fed at 7:00–8:00 a.m. and 6:00 p.m.

Pigs were randomly assigned to groups, ensuring they were matched by age, sex, and genotype when possible. Details on the number, ages, and sexes of animals are provided in **Supplementary Tables 3-6**. A random number generator was used for selection order randomization. No pigs received routine or consistent medical or pharmacological treatment for Angelman syndrome phenotypes; interventions were administered only when deemed necessary, such as under veterinary guidance or to address life-threatening conditions.

## METHOD DETAILS

### Comparative genomic and phylogenetic analyses

The genomic sequence of human *UBE3A* (hs1 chr15:23070379-23175611) was aligned against orthologous regions in distantly related species, including crab-eating macaque (*Macaca fascicularis*), house mouse (*Mus musculus*), Norway rat (*Rattus norvegicus*), domestic pig (*Sus scrofa*), and nine-banded armadillo (*Dasypus novemcinctus*) using Progressive Cactus^(59)^. To examine sequence conservation at the transcript and protein levels, *UBE3A* cDNA and amino acid alignments were generated for the same species using MUSCLE^(60)^. A maximum-likelihood phylogenetic tree was constructed from the cDNA sequences using IQ-TREE, with ultrafast bootstrap approximation for branch support^(61)^. The resulting phylogenetic tree was visualized using iTOL^(62)^. Conserved genomic regions were graphically represented using the Zoonomia alignment track^(63)^ within the UCSC Genome Browser (http://genome.ucsc.edu/).

### Generation of the UBE3A deletion pig model

Male porcine fetal fibroblast cells were isolated and transfected with a CRISPR plasmid (pX330-U6-Chimeric_BB-CBh-hSpCas9, Addgene #42230) using the Amaxa NHDF Nucleofactor Kit (Lonza, VPD-1001). Several CRISPR guide RNAs (**Supplementary Table 2**) were used. Transfected fibroblasts were isolated into single colonies, and PCR screening (**Supplementary Table 8**) was performed to identify guide RNAs generating a full deletion. CRISPRs 8 and 4 successfully produced a heterozygous deletion of *UBE3A*. Following confirmation of a CRISPR pair capable of generating the desired mutation, a second transfection was performed to isolate single colonies suitable as donor cells for somatic cell nuclear transfer (SCNT). SCNT was conducted as previously described^(64)^.

### Genotyping and Sanger sequencing

DNA was extracted from tissue samples using the Qiagen Blood and Tissue DNeasy Kit (Qiagen, 69504). Samples were incubated on a rocker at 56°C overnight for lysis. DNA was eluted in 200 µL of buffer AE, and concentrations were measured with a Qubit 4 Fluorometer using the Qubit dsDNA BR Assay Kit (Thermo Fisher Scientific, Q32850). Genotyping reactions were performed in a 20 µL total volume, containing 25–50 ng of DNA, 500 nM each of *UBE3A* 1-13F and *UBE3A* 1-13R2 primers, and 10 µL of 2× Phire Mix (Thermo Fisher Scientific, F126L) (**Supplementary Table 8**). Amplifications were carried out on a Bio-Rad T100 Thermal Cycler with the following conditions: an initial denaturation at 98°C for 30 seconds; 35 cycles of 98°C for 5 seconds, 64°C for 5 seconds, 72°C for 5 seconds; followed by a final extension at 72°C for 1 minute. PCR products were analyzed on an agarose gel, and target bands were excised for purification. Gel extraction was performed using the PureLink Quick Gel Extraction and PCR Purification Combo Kit (Thermo Fisher Scientific, K220001). The purified amplicons were sequenced using the same primers at the LGT Core at Texas A&M University.

### Genome sequencing

Genomic DNA from three pig samples (G-6 (father), 1-2 (mother), and 6-10 (offspring)) was sequenced using Illumina paired-end 150 base-pair reads at a depth exceeding 50x coverage. Raw sequencing reads were quality-filtered using Trimmomatic v0.39, and alignments were generated for each sample following the GATK v3 Best Practices Workflow^(65)^. Joint genotyping was also performed. Variant recalibration, posterior probability estimation, and variant effect prediction were conducted using the ENSEMBL SNP dataset (ENSEMBLE release 80) for pigs as the training set.

### CRISPR-Off analysis

Potential off-target sites for the sgRNAs (U3A-CRISPR8-Ex1-top and U3A-CRISPR4-Ex13-top) were predicted using CRISPOR^(66)^, identifying a total of 174 off-target intervals. These predicted intervals were cross-referenced with INDELs identified in previous analyses to assess potential off-target effects. To investigate donor plasmid insertion, unmapped reads from the maternal, paternal, and offspring samples were analyzed for the presence of the CRISPR plasmid (pX330-U6-Chimeric_BB-CBh-hSpCas9). A custom BLAST database was created to facilitate this search.

### RNA isolation and cDNA synthesis

RNA was isolated from tissues using the Qiagen RNeasy Plus Kit (Qiagen, 74136). Frozen tissues were weighed, lysed in buffer, and homogenized with a TissueLyser II (Qiagen, 85300). RNA purification followed the manufacturer’s protocol, and RNA concentration was measured using a Qubit 4 Fluorometer with the Qubit RNA BR Assay Kit (Thermo Fisher Scientific, Q10211). For cDNA synthesis, 200 ng of RNA was reverse-transcribed in a 50 µL reaction using the High-Capacity RNA-to-cDNA Kit (Thermo Fisher Scientific, 4388950).

### qRT-PCR

#### UBE3A

TaqMan assays were performed in a 10 µL reaction volume containing 2 µL of cDNA, 1× Gene Expression Master Mix (Thermo Fisher Scientific, 4369016), and 1× TaqMan Gene Expression Assay (Thermo Fisher Scientific). The reference assay (PPIA, Ss03394782_g1) and target assay (UBE3A, Ss04323425_m1) were duplexed in each reaction. ARFGAP2 (Ss04327825_m1) was used as the reference in retina and was single-plex. Cycling conditions included 2 minutes at 50°C, 10 minutes at 95°C, and 40 cycles of 15 seconds at 95°C and 1 minute at 60°C, with fluorescence measured during the 60°C step.

#### UBE3A-AS

Total reaction volume was 10ul, including 2ul of cDNA, 1X PowerUp SYBR Green Master mix (ThermoFisher Cat# A25741), and 500nM of each primer (forward and reverse). Primers were UBE3A-AS (F: CTGAATGGGACCTGCTGTCT R: TTTTGTAGTTTTGACCTCACATGC) and PPIA (F:CATGGTTAACCCCACCGTCT R: TGCAAACAGCTCGAAGGAGA). Cycling conditions were 2 minutes at 50C, 2 minutes at 95C, and 40 cycles of 15 seconds at 95C and 1 minute at 60C, with readings taken at the 60C step of every cycle. Quantitative RT-PCR reactions were run on a Bio-Rad CFX96 Touch Real-Time PCR Detection System, and data were analyzed using CFX Maestro software. Samples with PPIA Ct values >30 were excluded. Data quality was visually inspected for discrepancies between technical replicates and potential plate effects.

### Western blot

Tissue samples were lysed in NP40 buffer (1% Nonidet P40, 0.01% SDS) with protease inhibitors (Roche, 11836153001). Lysates were mixed with 4× Laemmli buffer (Bio-Rad, 1610737) containing β-mercaptoethanol and heated at 95°C for 10 minutes. Proteins were separated on a 10% Mini-PROTEAN TGX Stain-Free Gel (Bio-Rad, 4568033) at 30 V for 30 minutes, then 100 V for 45 minutes, and transferred to a nitrocellulose membrane using the Trans-Blot Turbo System (Bio-Rad, 1704150). The membrane was blocked with 5% non-fat dry milk in TBST (Tris-buffered saline with 0.1% Tween 20) for 1 hour at room temperature, then incubated overnight at 4°C with mouse anti-UBE3A antibody (1:1000; BD Biosciences, 611416). After washing, the membrane was incubated with goat anti-mouse HRP-conjugated secondary antibody (1:1000; Invitrogen, 626520) for 1 hour at room temperature. Protein detection was performed using Clarity Western ECL Reagent (Bio-Rad, 1705060) and visualized on a ChemiDoc Imaging System (Bio-Rad). Protein levels were quantified and normalized to the amount of total protein per sample using the stain-free image.

### Immunostaining

Tissues were sectioned into 4 mm coronal slices and fixed in 10% neutral buffered formalin (NBF). Tissue processing and staining were conducted by the Texas A&M University Veterinary Medicine and Biomedical Sciences Core Histology Laboratory. Coronal slices were paraffin-embedded and sectioned into 4.5 µm slides. DAB (3,3′-diaminobenzidine) immunostaining was performed using the UBE3A antibody (Sigma-Aldrich, SAB1404508) at a 1:750 dilution, with all slides processed under identical conditions, followed by hematoxylin counterstaining. Slides were scanned at 40× magnification using a 3DHistech Pannoramic Scan II.

### Capture of Characteristic Phenotypes

Animals were video recorded and observed in their home environments under standard housing conditions to document characteristic home change behaviors, which minimized stress and captured natural behaviors.

### Muscle tone analysis

A group of trained, blinded observers together assessed muscle tone on a 1–5 scale, where 1 indicated very low muscle tone with prominent shoulders, spine, and hips, to 5 denoting well-muscled pigs with no visible ribs or hip bones.

### Modified acute restraint stress test

Three groups of pigs (ages 1–58 days old) were temporarily restrained multiple times. Struggling was scored on a 1–3 scale: 1 for no movement, 2 for leg kicking, and 3 for whole-body thrashing. Audible vocalization was also scored on a 1–3 scale: 1 for no vocalization, 2 for low grunts, and 3 for squealing.

### Incline test

A mobile pen with a slatted floor was modified to include an approximately 30-degree slope. The pen was disinfected, rinsed, and dried before each use to prevent slipping. Each pig was tested twice in different starting positions—once facing upward and once facing downward—to assess navigation ability. Trials lasted up to 1 minute or until all four hooves reached the flat section. Successful navigation was defined as controlled descent without stumbling or falling. Pigs that were stationary for the full minute were excluded from analysis.

### Righting-reflex

Pigs were placed in a supine position and the time taken to self-right and stand on all four hooves was recorded, as previously decided for many species.

### Open field arena specifications and Vocalization in an open field environment

The open field arena measured 10 ft × 10 ft, enclosed by 4 ft matte black walls for video tracking, with a floor of commercial black stall mats. A ceiling-mounted camera centered above the arena and a microphone on the top left-hand wall captured data. The arena was cleaned with Rescue disinfectant (Virox Technologies Inc, #:23305), rinsed with water, and dried between animals. Auditory vocalizations were recorded during a 6-minute open field test using a microphone (Zoom H1n, B07J5M244T) positioned on the arena’s top left-hand wall. Audio files were analyzed with Noldus UltraVox software to generate spectrograms. A trained, blinded observer manually counted vocalizations while reviewing the spectrogram. Vocalizations were categorized by frequency: 0–8000 Hz as “low grunts,” 8001–12,500 Hz as “high grunts,” and 12,501–22,000 Hz as “squeals”.

### Gait analysis

The TekScan Strideway system was used to evaluate pig gait. Pigs were acclimated to the handler for 5– 10 minutes daily over three days in their home pen, followed by two days of training to walk across the system using treats for compliance. Gait was recorded once daily for three consecutive days, yielding three recordings per pig. The system was calibrated daily before use and disinfected between animals. Pigs unable to produce three consecutive steps per hoof or exhibiting destructive or uncooperative behavior were excluded. A trained observer selected continuous steps suitable for analysis, requiring at least 12 steps (three consecutive steps per hoof) without pauses. TekScan Strideway software was used to calculate average step width and length.

### Assessment of open field locomotion

Pigs were individually placed into the open field arena, given a 1-minute acclimation, and then recorded for 5 minutes. Locomotion was quantified using Noldus EthoVision software, measuring the percentage of time in motion, distance traveled and velocity. Movement was defined as a bodily position change exceeding 30 cm/s (0.68 mph), non-locomotive behaviors, such as head shaking and rooting, were excluded.

### Body weight

Pigs were weighed at multiple time points during development and categorized into age groups to account for variations in weighing times. Body weights were compared among genotypes within each age group to assess potential differences.

### Gross brain weights

Day 10 and 65 pigs were anesthetized, perfused with PBS, and exsanguinated before brain removal and weighing. Day 130 pigs were euthanized without perfusion prior to brain removal and weighing. Brain weights were compared among genotypes in age-matched pigs to assess potential differences.

### Magnetic resonance imaging (MRI)

All pigs underwent MRI scans using a 3T scanner (Magnetom Verio, Siemens Healthineers, Erlangen, Germany). Subjects were positioned head-first-prone and the 16-channel body array anterior coil and the spine coil were used. Three-dimensional T1-weighted gradient echo (MP-RAGE) images were acquired in the sagittal plane with the following parameters: repetition time of 2100 ms, echo time of 2.64 ms, field-of-view of 200 mm with 100% phase, matrix size of 256 mm x 256 mm, 0.98 mm slice thickness, and a 12-degree flip angle. Brain volumes were estimated by manual segmentation using Siemens Inveon Research Workplace (IRW) software, version 4.1.

### Quantification and statistical analysis

The sample size was determined based on available funding and logistical feasibility rather than a formal statistical power calculation. While this approach may limit the statistical power of our findings, we ensured that the number of animals used was sufficient to provide meaningful biological insights while adhering to ethical guidelines for animal research. All statistical analyses were performed using GraphPad Prism and JMP Pro 16. Statistical significance was set at p < 0.05. Prior to performing statistical analyses, sex differences were assessed in studies with a sufficient number of animals. No significant sex differences were observed in any studies; therefore, data were not stratified by sex. For sample sizes less than 10, individual data points were displayed on graphs. Binary outcomes (e.g., incline test) were compared using two-tailed Fisher’s exact tests. Ordinal data (e.g., muscle tone) were analyzed using Chi-square ordinal logistic regression or ordinal logistic regression, with false discovery rate correction applied where appropriate. Continuous variables (e.g., righting reflex, and brain weights) were analyzed using Wilcoxon two-sample exact tests or Student’s t-tests, with ANOVA and Tukey-Kramer HSD post hoc tests for multiple group comparisons. Neuroimaging data (brain MRI measurements) were analyzed using Student’s t-tests. Data normality was assessed using the Shapiro-Wilk test, and parametric or non-parametric tests were applied accordingly.

## ACKNOWLEDGEMENTS

We thank the following individuals for their contributions to the project: John Griffin, James Elliot, Clay Ashley, Jennifer Fridley, Destiny Taylor, Richard Colson, Hillary Shaheen, Ruth Robinson, Kamelia Radeva, Niamh Nichols, Julie Gonzales, Tara Woeste, Dieter Ally, Desiree Slaughter, Aubrey Rinderknecht, Katie Bumgardner, Charles Collins, Joseph Ready, and Brittany Leatherwood. We also extend our thanks to the staff at the Veterinary Medical Park and the Texas Institute for Preclinical Studies at Texas A&M University. The authors would like to thank the Texas A&M Institute for Genome Sciences and Society (TIGSS) for providing computational resources and systems administration support via the TIGSS High-Performance Computing Cluster (TIGSS-HPC). We are also grateful to the Translational Imaging Center at the Texas A&M Institute for Preclinical Studies for providing imaging resources.

## AUTHOR CONTRIBUTIONS

Conceptualization was performed by LSM, SGC, FIRE, JP, and SVD. Data curation was conducted by LSM, SGC, SS, RS, KK, and SVD. Formal analysis was carried out by LSM, TJ, and SVD. Funding acquisition was contributed by SVD. Investigation was conducted by LSM, SGC, CT, LM, TJ, DR, LS, WF, OH, MH, AT, AC, AS, BR, CK, CC, WJM, and SVD. Methodology was developed by LSM, SGC, SS, RS, CT, WF, JP, and SVD. Project administration was handled by SGC, CT, and JP. Resources were provided by WF, and JP. Supervision was conducted by SGC, JP, and SVD. Validation was carried out by SVD. Visualization was performed by LSM, SGC, TJ, AT, and SVD. Writing—original draft preparation was done by LSM, SGC, and SVD. Writing—review and editing were conducted by LSM, SGC, and SVD.

