## Supplementary Figure for "A preclinical pig model of Angelman syndrome mirrors the early developmental trajectory of the human condition"

### SUPPLEMENTARY FIGURES

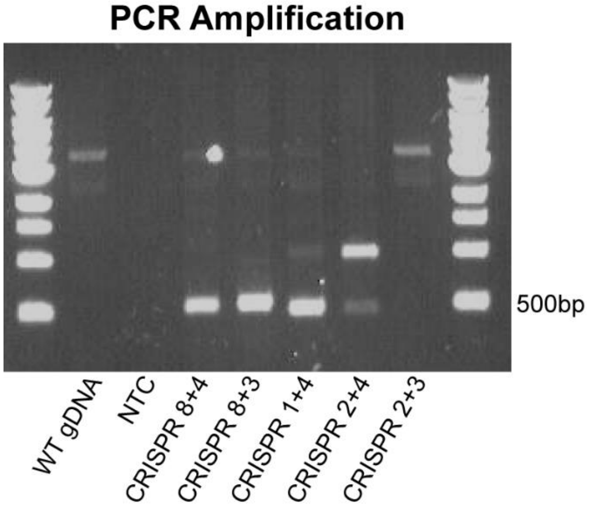


**Supplementary Figure 1. PCR validation of CRISPR-mediated deletion in fibroblast colonies.** Agarose gel showing PCR amplification of genomic DNA from fibroblast colonies transfected with various CRISPR guide RNA combinations. Lanes include wild-type (WT) genomic DNA and no-template control (NTC).

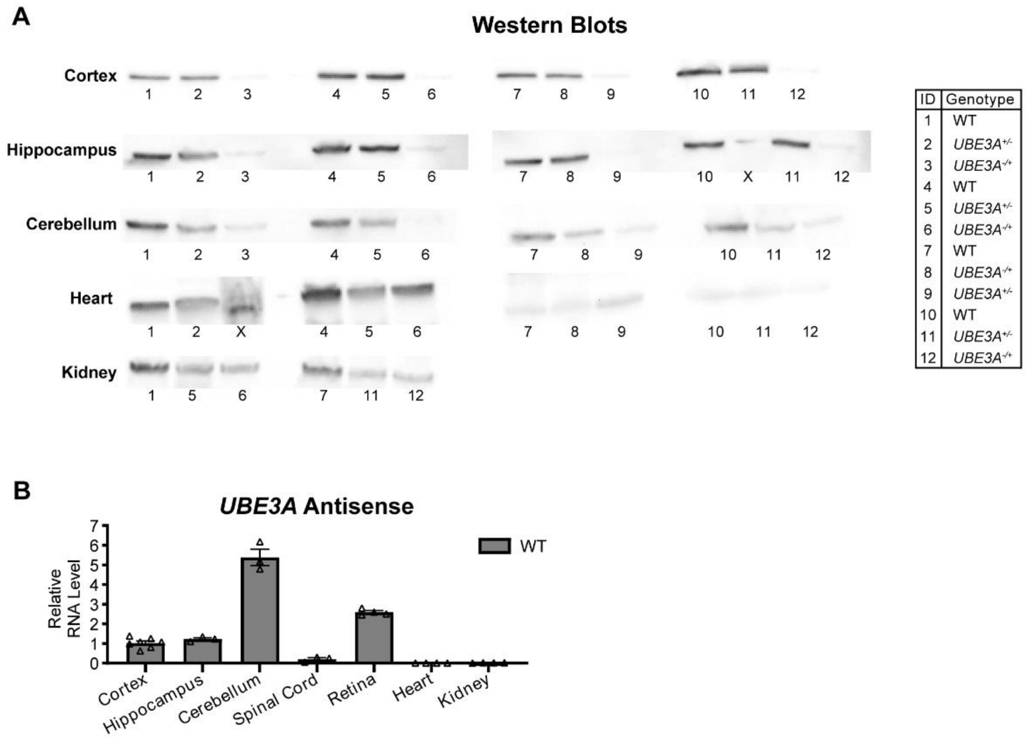


**Supplementary Figure 2. Western blots and pig *UBE3A-AS* transcript is expression.** (**A**) Western blots showing UBE3A protein expression across different tissues and genotypes. ‘X’ denotes wells not used in the analysis. (**B**) RT-PCR quantification of *UBE3A-AS* expression in WT tissues, normalized to cortex. Data presented as mean ± SEM.

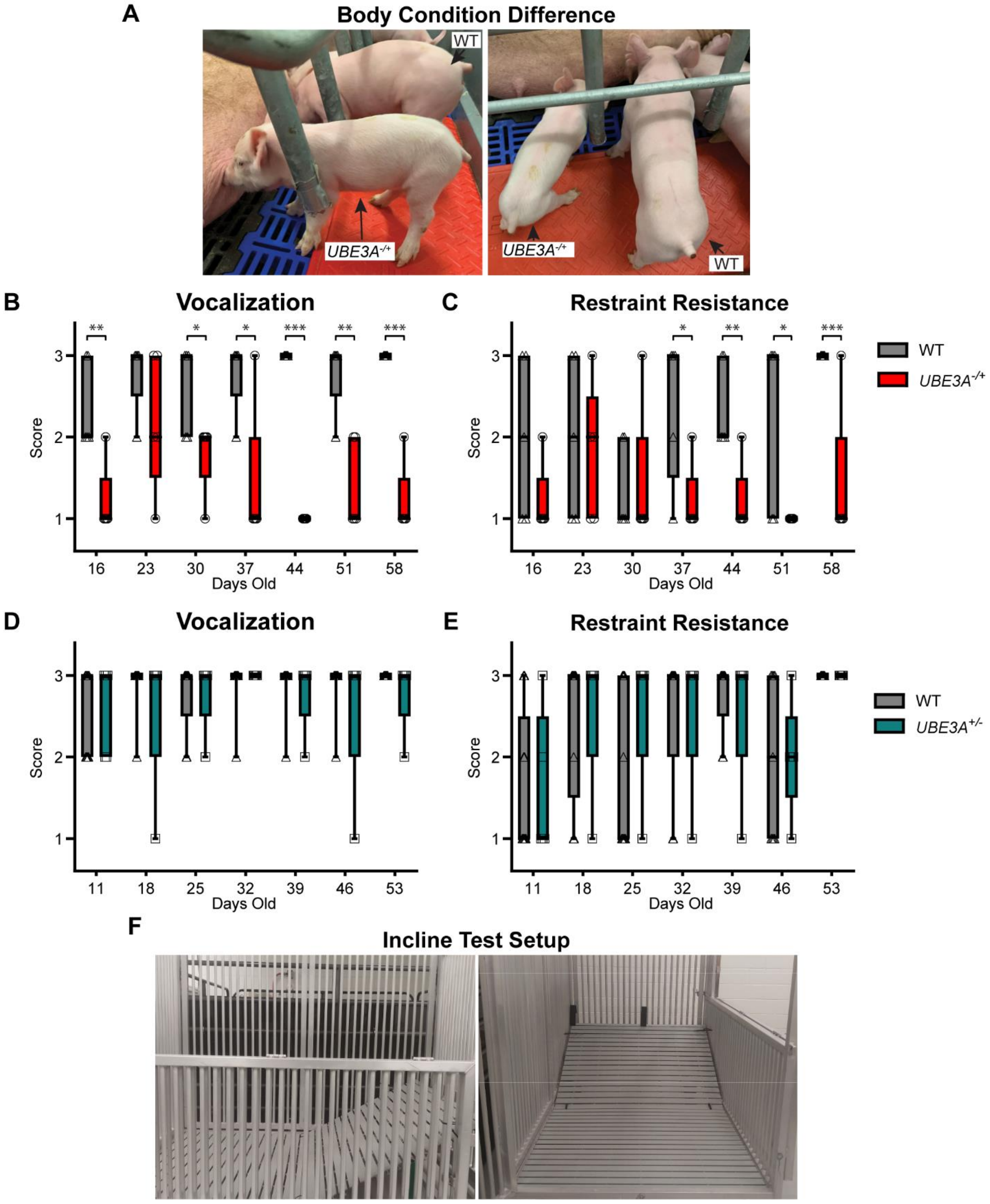


**Supplementary Figure 3.** ***UBE3A****$^{-/+}$* **pigs have impaired neonatal and adolescent development.** (**A**) Representative images showing size and muscle tone differences between *UBE3A$^{-/+}$* and WT pigs. (**B and C**) *UBE3A$^{-/+}$* pigs had significantly lower vocalization and resistance scores during restraint tests compared to WT pigs. Data distribution presented as a boxplot, Ordinal Logistic Fit, FDR correction. (**D and E**)

*UBE3A*$^{+/-}$ pigs had similar vocalization and resistance scores during restraint tests compared to WT pigs. Data distribution presented as a boxplot, Ordinal Logistic Fit, FDR correction. Vocalization scores = 1: no vocalization; 2: grunting; 3: squealing, and resistance scores = 1: no movement; 2: leg kicking; 3: whole-body thrashing. (**F**) Custom slope used for incline and decline testing. Abbreviations: WT = wild-type, *UBE3A*$^{-/+}$ = maternal *UBE3A* deletion and *UBE3A*$^{+/-}$ = paternal *UBE3A* deletion. *$P$ < 0.05, **$P$ < 0.01, and ***$P$ < 0.0001.

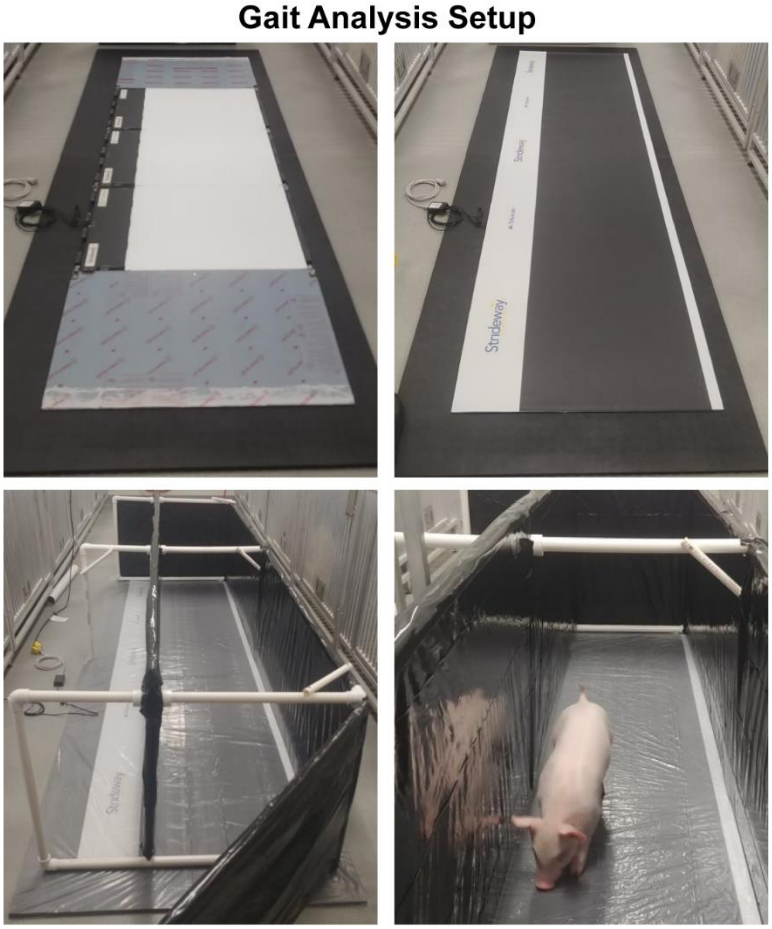


**Supplementary Figure 4. Gait analysis setup using the TekScan Strideway System.** Images of the TekScan Strideway gait mat and custom-made chute used to guide pigs during gait analysis. The chute ensured straight walking along the gait mat and prevented turning, enabling accurate measurement of gait parameters, including step width, stride length, and walking patterns.

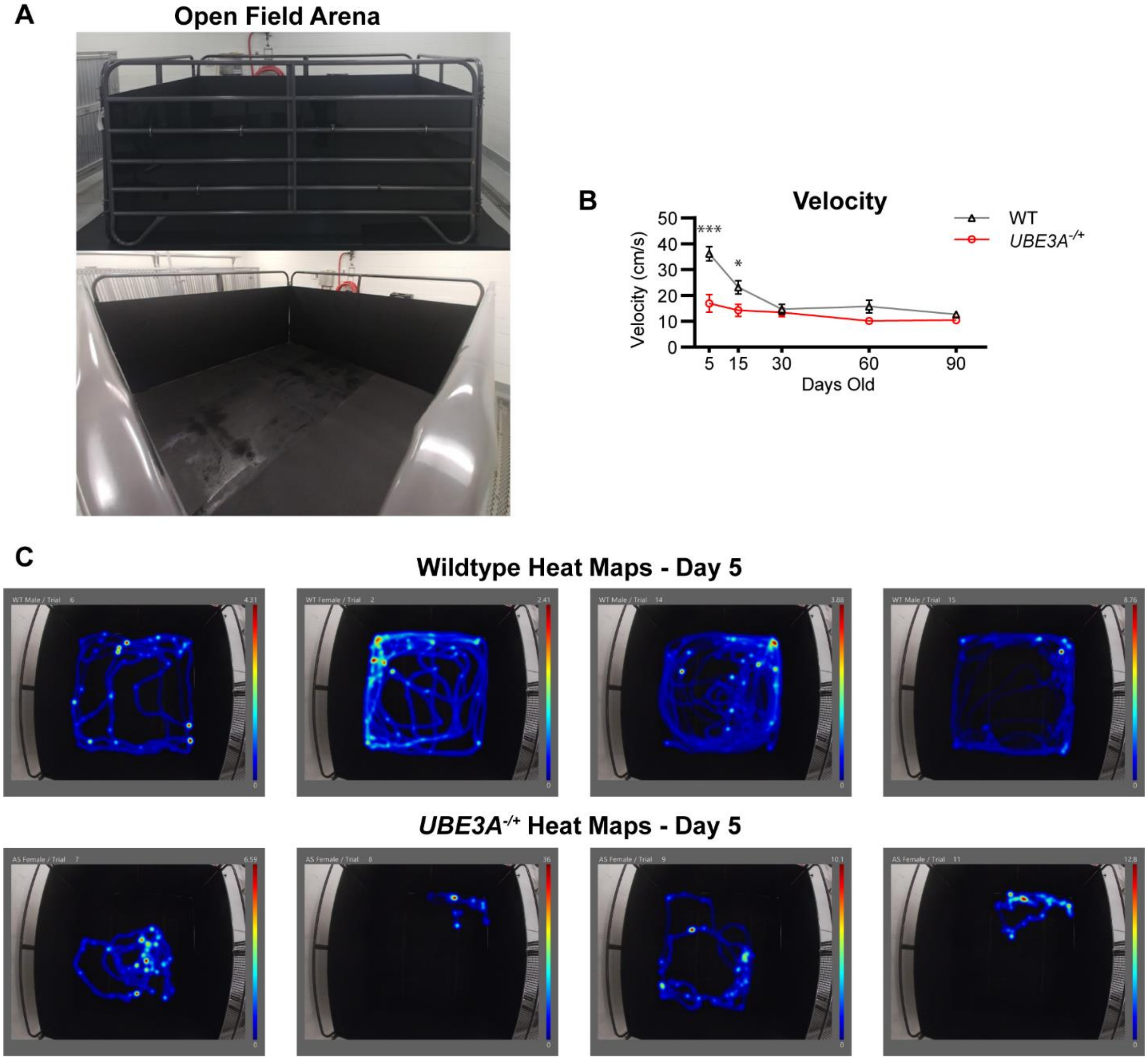


**Supplementary Figure 5. Open field arena for pigs and *UBE3A*$^{-/+}$ pigs demonstrate hypoactivity.** (**A**) Open field arena (10 ft × 10 ft) used to assess pig activity levels. (**B**) Velocity measurements show significantly reduced movement in *UBE3A*$^{-/+}$ pigs compared to WT littermates at 5 and 15 days old. Data presented as mean ± SEM, Students t, all pairwise comparison. (**C**) Representative heat maps showing the movement of WT and *UBE3A*$^{-/+}$ pigs in the open field arena. Abbreviations: WT = wild-type and *UBE3A*$^{-/+}$ = maternal *UBE3A* deletion. $*P < 0.05$, $**P < 0.01$, and $***P < 0.0001$.
